# Human posterior parietal cortex responds to visual stimuli as early as peristriate occipital cortex

**DOI:** 10.1101/377887

**Authors:** Tamar I. Regev, Jonathan Winawer, Edden M. Gerber, Robert T. Knight, Leon Y. Deouell

**Affiliations:** Edmond and Lily Safra Center for Brain Science, Hebrew University of Jerusalem, Jerusalem, Israel; Department of Psychology, New York University, New York, New York, USA; Helen Wills Neuroscience Institute, University of California, Berkeley, California, USA; Department of Psychology, Hebrew University of Jerusalem, Jerusalem, Israel

**Keywords:** Early visual processing, onset latency estimation, electrocorticography, ECoG

## Abstract

Much of what is known about the timing of visual processing in the brain is inferred from intracranial studies in monkeys, with human data limited to mainly non-invasive methods with lower spatial resolution. Here, we estimated visual onset latencies from electrocorticographic (ECoG) recordings in a patient who was implanted with 112 sub-dural electrodes, distributed across the posterior cortex of the right hemisphere, for pre-surgical evaluation of intractable epilepsy. Functional MRI prior to surgery was used to determine boundaries of visual areas. The patient was presented with images of objects from several categories. Event Related Potentials (ERPs) were calculated across all categories excluding targets, and statistically reliable onset latencies were determined using a bootstrapping procedure over the single trial baseline activity in individual electrodes. The distribution of onset latencies broadly reflected the known hierarchy of visual areas, with the earliest cortical responses in primary visual cortex, and higher areas showing later responses. A clear exception to this pattern was robust, statistically reliable and spatially localized, very early responses on the bank of the posterior intra-parietal sulcus (IPS). The response in the IPS started nearly simultaneously with responses detected in peristriate visual areas, around 60 milliseconds post-stimulus onset. Our results support the notion of early visual processing in the posterior parietal lobe, not respecting traditional hierarchies, and give direct evidence for the upper limit of onset times of visual responses across the human cortex.

## Introduction

Measuring the timing of neural responses in distinct cortical areas is important for understanding the network dynamics sub-serving sensory information processing in the brain. In particular, estimation of relative onset latencies can be used to test hierarchical relations and cortical connectivity. Specifically in the visual system, onset latencies of responses at various cortical and sub-cortical structures have been used to map functional connectivity and distinguish streams of processing (Schmolesky *et al.*, 1998; Schroeder *et al.*, 1998; Bullier, 2001; Chen *et al.*, 2007; Ledberg *et al.*, 2007). However, such measurements have been done primarily in laboratory animals.

Due to the difficulty of measuring human brain signals with both high spatial and temporal resolution less is known about the precise spatio-temporal evolution of visual information processing in the human brain. In fMRI studies, the spatial resolution allows functional localization of specific parts of the visual system (such as V1, V2 etc.) using retinotopic mapping (e.g., Dumoulin & Wandell, 2008; Wandell & Winawer, 2011), but the temporal resolution is too low to study the latency of neural responses. Scalp EEG studies give good indication for the timing of processing (e.g. Foxe & Simpson, 2002). However, electrical measures on the scalp sum over an unknown number anatomical sources, resulting in poor spatial resolution, as well as temporal smearing. Together with the ill-posed inverse problem of EEG source reconstruction, decomposing the EEG signal into its constituent components and localizing each component is a major challenge. MEG is similarly constrained by the inverse problem and spatial summation, but since magnetic fields are less distorted by the conductive medium than electric fields, localization is arguably more accurate than with EEG, while maintaining the same high temporal resolution. EEG and MEG studies have shown considerable variability in estimation of the onset time of visual responses (see summary in Shigihara *et al.*, 2016). Some reported surprisingly early onsets in occipital cortex, as early as 25-30 ms (Moradi *et al.*, 2003; Inui & Ryusuke, 2006; Inui *et al.*, 2006; Shigihara & Zeki, 2013; 2014), or in some cases even at 10-15ms post onset (Shigihara *et al.*, 2016). As previously argued (Shigihara *et al.*, 2016), it is hard to compare absolute onset times between studies because onset times are affected significantly by the luminance, contrast, spatial frequency, size, and location in the visual field of the stimuli. Rather, the relative latencies are most directly interpretable. The results of studies by Zeki and colleagues (ffytche *et al.*, 1995; Shigihara & Zeki, 2013; 2014; Shigihara *et al.*, 2016), suggested parallel activation of V1 and peristriate cortex (i.e. with no lag), although due to the limitation of the method the precise location of the early activity could not be determined.

Here we present an estimation of onset latencies of visually evoked responses across the posterior human cortex using electrocorticographic (ECoG) recordings, having the advantage of both spatial and temporal high resolution, in a single human patient requiring surgery for intractable epilepsy. The patient went through pre-surgical evaluation that included functional MRI followed by implantation of sub-dural electrodes covering various posterior cortical areas. We examined the spatio-temporal progression of visual responses in the cortex and examined the degree to which they abide by the established streams of processing. For that aim, we developed a statistical method for assessment of onset latencies in continuous signals using bootstrapping of baseline activity.

## Materials and Methods

### Patient

The patient, who suffered from intractable epilepsy, was hospitalized for pre-surgical evaluation at the Stanford Medical Center. As part of this clinical procedure, the patient was implanted with sub-dural electrodes, to allow a more precise localization of the epileptic focus, so that it can be subsequently surgically removed. The number and location of the electrodes was determined solely based on clinical needs, and, before electrode implantation, the patient signed an informed consent to participate in our experiment. All procedures performed in this study were approved by the UC Berkeley Committee on Human Research and corresponding IRBs at the clinical recording site. The patient was a right handed, 45 years old male, English speaking, with normal intellectual abilities and no psychiatric or visuospatial abnormalities, as revealed by standard pre-surgical evaluation.

### Stimuli

The stimuli consisted of grayscale pictures of four categories; human faces, watch faces, drawings of everyday objects, and pieces of clothing, which were the target stimuli. The categories of visual images served the purpose of another experiment and are largely ignored in the present study focusing on early visual areas. The stimuli were 5.4cm square on the background of a black 34.4×19.3cm LCD screen (refresh rate 60Hz), located about 65cm away from the subject’s eyes. Hence, the stimuli subtended about 4.6 degree of visual angle, in the center of the visual field. The experiment was run under normal hospital room lighting.

### Procedure

The patient was half seated in his hospital bed with the laptop computer used to present the stimuli placed on a table suspended above (but not touching) his legs. The experiment was controlled by Eprime 2 running on Windows XP (Microsoft, Inc.) operating system. The position of the laptop allowed the patient to press the space key on the keyboard to indicate target detection. A photodiode on the bottom screen corner detected a bright rectangle which was presented on the screen together with the image. The bright rectangle was completely covered by the photodiode unit so that it was invisible to the subject. The signal from the photodiode was recorded alongside the EEG and later used to create stimulus triggers. The identity of each stimulus was recorded by Eprime and later merged with the triggers extracted from the photodiode channel. The lag between the onset of stimulus in the middle of the screen and that of the rectangle in the bottom-right corner of the screen was 5 ms, as measured using an analog oscilloscope after the experiment. This delay was taken into account in the analysis.

During the experiment, the patient was required to press the space key whenever he detected a piece of clothing (nearly 10% of stimuli). The experiment consisted of 8 blocks of 86 visual images mixed across the non-target categories with equal shares. The duration of stimuli was 300, 600, 900, 1200 or 1500 milliseconds (ms), with equal probabilities. The variable durations served the purpose of another experiment and are largely ignored in the present study focusing on onset responses (Gerber *et al.*, 2017). The order of the stimuli and durations were quasi-random. Inter-stimulus intervals (ISI) were between 600, 900, 1200 or 1500 ms, randomly distributed. Before the experiment started the patient was presented with a sample of the targets, and with a few practice trials, which were not recorded. In total, 688 stimuli were presented during the experiment.

### ECoG recording

The patient was implanted with 112 sub-dural electrodes covering areas of the occipital, parietal and temporal cortices of his right hemisphere (Fig. 2). The electrodes (AdTech Medical Instrument Corp.) were 2.3-mm in diameter, and arranged in either 2D grids or 1D strips (Fig. 2). Neighboring electrodes were approximately 0.5-1 cm center-to-center apart from each other within a single grid or strip. The EEG signal was recorded with a Tucker-Davis Technologies (TDT) recording system at a sampling rate of 3051.76 Hz, with an online 0.5 Hz high-pass filter.

**Figure 1.**
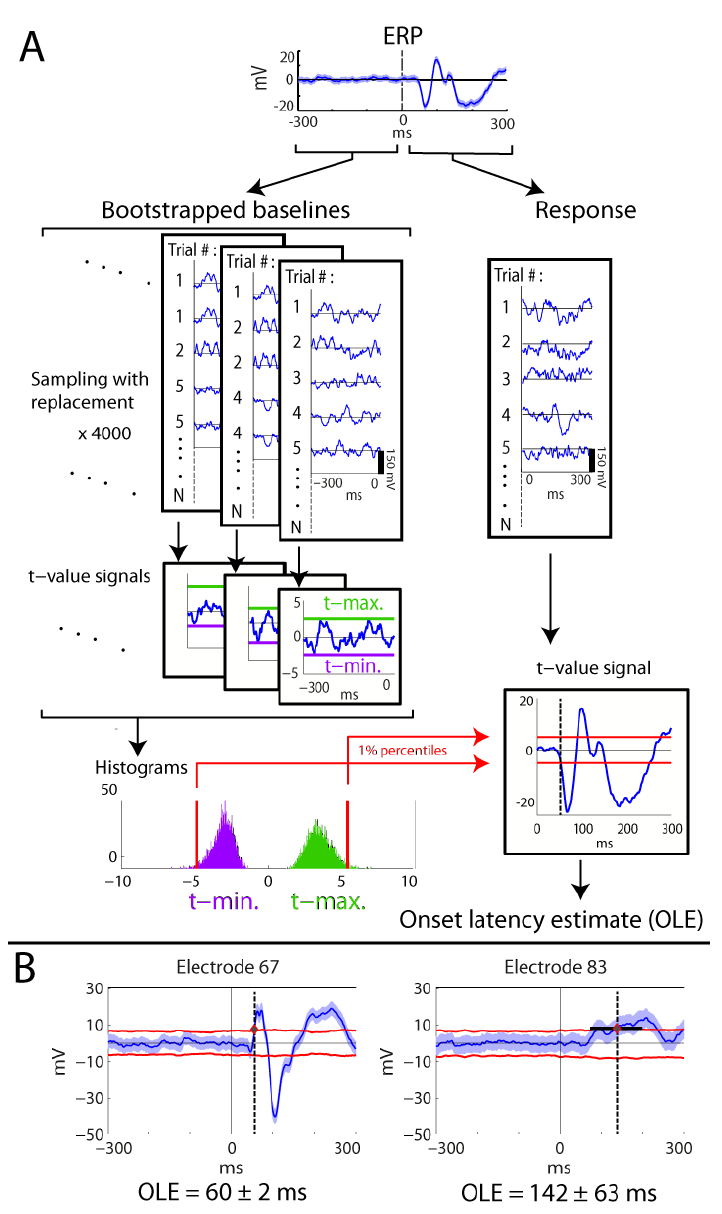
Illustration of the onset latency estimation method. **A.** Calculation of onset latency estimate (OLE) for a specific electrode via bootstrapping over the baseline trials, construction of empirical distributions of maximal and minimal tvalues, and using the 0.01 percentile values as thresholds for the response t-value signal (see methods). **B.** Estimation of temporal error of the OLE. Examples of specific electrodes having small or large temporal errors (left and right, respectively). Shaded blue area is 99% confidence interval around the mean. Red lines are the translation of the constant t-value thresholds back to the voltage domain, multiplying by the instantaneous standard error (see methods for detailed explanation). Electrode numbers are specified in Fig. 2.

**Figure 2.**
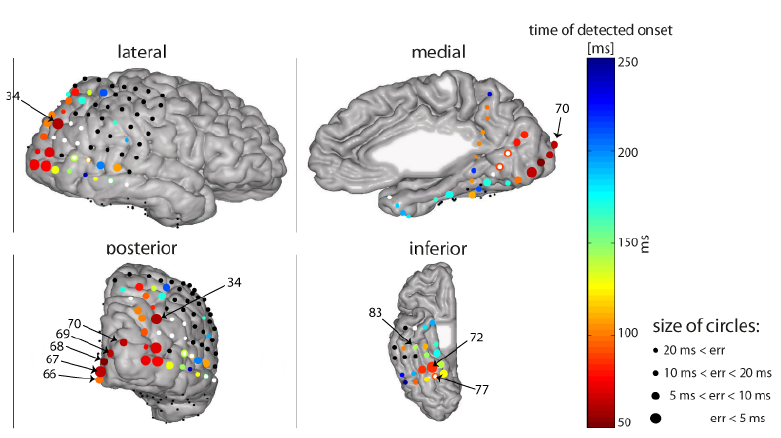
Significant visual onset latencies in all electrodes. Four views of the same brain are presented. The color of the circles denotes the timing of the estimated onset (see color bar). The size of the circles denotes the temporal error estimate (see methods): larger circles indicate more reliable estimates (smaller temporal errors; see legend). White dots indicate electrodes that were labeled as epileptic or with other electric artifacts. However, we looked for onset latencies of the bad electrodes as well, and if a significant onset was found, the colored circle was placed under the white dot (e.g. electrode 78). Black dots indicate electrodes for which no significant onset was detected. Numbers are indicated here for specific characteristic electrodes, and electrodes that are referred to in the text. Note that some electrodes appear in more than one view, however their onsets and/or numbers might appear in some of the views, e.g., the onsets of inferior electrodes are not indicated in the posterior view. Reference: common average (CAR). Electrodes that are explicitly mentioned in the text are labeled on the posterior and inferior views. For a dynamic depiction of activation over time see supporting information video S1.

### Electrode localization

Electrode locations were identified manually using BioImageSuite (www.bioimagesuite.org) on a post-operative Computed Tomography (CT) scan coregistered to a pre-operative MR scan using the FSL software package (Jenkinson & Smith, 2001; Jenkinson *et al.*, 2002). Individual subjects’ brain images were skull-stripped and segmented using FreeSurfer (http://surfer.nmr.mgh.harvard.edu). Localization errors driven by both co-registration error and anatomical mismatch between pre- and post-operative images were reduced using a custom procedure which uses a gradient descent algorithm to jointly minimize the squared distance between all electrodes within a single electrode array/strip and the cortical pial surface (see Dykstra *et al.*, 2012 for a similar procedure). In contrast to methods that only attempt to correct individual electrodes’ position in relation to the pial surface, this method preserves the original array topography, thus providing a more reliable estimate of actual electrode positions. A related method (Hermes *et al.*, 2010) was shown to localize electrodes to within 2-4 mm based on ground truth measures from intraoperative photography. We report the MNI coordinates of electrode 34 (see Results section ‘Very early visual response at posterior intra-parietal sulcus’), based on surface registration to an MNI152 standard-space T1-weighted average structural template image.

### Co-registration with retinotopic mapping

Retinotopic mapping was performed pre-operatively, as described in detail in a prior publication involving the same patient (Winawer *et al.*, 2013, sections ‘Stimuli for fMRI Experiments’ and ‘Anatomical and Functional MRI’). In brief, the subject was presented with a drifting checkerboard contrast pattern revealed within a moving bar aperture during nine 3.5-minute scans. The functional data were co-registered to a whole-brain T1-weighted image (1×1×1 mm), which was segmented into gray and white matter using Freesurfer’s autosegmentation tool. The autosegmentation produces a cortical ribbon for each hemisphere, and the functional time series were resampled to the cortical ribbon via trilinear interpolation. Population receptive field models were solved on this resampled data, and model parameters – polar angle and eccentricity – were visualized on the cortical surface. Retinotopic maps were identified on the cortical surface following previous definitions of visual field maps and their boundaries (Winawer *et al.*, 2010; Wang *et al.*, 2014).

### Data analysis

The analysis was performed using Matlab software (The Mathworks Inc., Natick, MA) versions 2013b or 2016b, (yielding the same results) running on a Windows-based computer. The analysis steps are listed below.

### Pre-processing

Ten electrodes that were diagnosed clinically by the neurologist (Dr. Josef Parvizi) as including epileptic activity were ignored. The ongoing data was further reviewed by authors RTK and LYD and 6 more electrodes were ignored due to repeated electrical artifacts. For the remaining electrodes, temporal periods in which epileptic activity or other occasional artifacts were observed anywhere were excluded from the analysis across all electrodes.

The data were downsampled to 1000 Hz, and the 60Hz oscillation caused by US electricity system was filtered out using a notch filter version that was developed in our lab (Keren *et al.*, 2010), which mainly removes the ongoing oscillatory component, with less effect on transients. We used Common Average Reference (CAR) – a point-by-point average of all electrodes except for the epileptic and artifact-laden ones, or the Current Source Density (CSD) local reference, as explained below in section ‘Current Source Density (CSD) estimation’.

### Event Related Potentials (ERPs)

Since we were interested in the onsets of early visual responses irrespective of visual category or stimulus duration, the data was segmented to epochs lasting 600 ms, starting 300 ms before and ending 300 ms after stimulus onset. The baseline mean of the 300 ms preceding stimulus onset in each trial was subtracted from each time point of the segment, and all non-target trials (excluding artifacts) were averaged to calculate Event Related Potentials (ERPs) per electrode. The ERPs thus consisted of 569 trials.

### Onset latency estimation

To determine the first time point in which the ERPs significantly depart from baseline activity, we calculated the point-by-point one-sample t-value (against zero) across the trials per each electrode - 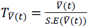 whereas 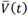 is the instantaneous average voltage across trials, and 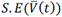 is the instantaneous standard error across trials. To determine a threshold t-value, we used a bootstrapping procedure over the 300 ms pre-stimulus baseline trials, which provided a distribution of expected t values under the null hypothesis of no response (Fig. 1.a). The bootstrapping procedure consisted of generating 4000 different groups of n=569 trials, by sampling with replacement from the original group of n baseline trials (see Fig. 1.a). The number 4000 was determined such that increasing it did not change the final onsets, up to 1ms. For each group of n bootstrapped trials, a baseline surrogate t-statistic signal (against 0, see formula above) was calculated point-by-point during the −300 to 0 pre-stimulus periods, and the maximal and minimal values of the t-statistic signal were noted. Histograms of the frequency distribution of maximal and minimal surrogate t-statistic values were generated for each electrode and the right and left 1% percentiles of the distribution were used as negative and positive thresholds for the actual response tstatistic signal, respectively (see Fig. 1A). The onset latency estimate (OLE) for each electrode was determined as the first time-point at which the true t-value signal of the response (0-300 ms) passed either the high or the low threshold t-values for that electrode. Consequently, the equivalent alpha level was 0.02. Next, in order to compute a temporal error for the OLE, the t-value thresholds were converted back to voltage by multiplication of the constant t-value threshold by the instantaneous standard error of the voltage at each time point of the response trials. The temporal error margin was determined as the interval surrounding the OLE, in which the voltage threshold was within the 99% CI around the ERP mean (i.e. the threshold was smaller than ERP + CI/2 or larger than ERP – C/2). The temporal error was calculated as half of the length of this interval (see Fig. 1B). Fig. 1 illustrates the onset latency estimation procedure.

### Current Source Density (CSD) estimation

Due to volume conduction, the activity measured in a given electrode may reflect neural activity that is remote from the location of the electrode. To mitigate the effect of volume conduction, we also used CSD to estimate onset latencies (Carvalhaes & de Barros, 2015), which effectively applies a high-pass spatial filter to the voltage measurements. Due to the nature of volume conduction, activity conducted from far away sources should be quite similarly measured by adjacent electrodes. In contrast, local sources will be measured much more strongly by the electrode near the source, and the measured activity will drop sharply in nearby electrodes. The current source density map thus filters out widespread activity, which is suspect of being conducted from remote sources, and highlights local current sources.

The CSD is inversely proportional to the Laplacian of the voltage field: *CSD* = –∇^2^*V*. It can be estimated easily by subtracting a local reference for each electrode (Hjorth, 1975; Schroeder *et al.*, 1998; Butler *et al.*, 2011). Effectively, the average voltage of the 4 or 2 neighboring electrodes is subtracted for each electrode residing on a 2D grid or 1D strip, respectively:

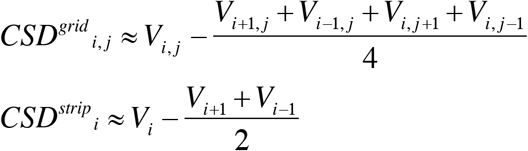

Notably, we did use electrodes that were marked epileptic as reference electrodes if they were neighboring other electrodes of interest. However, we excluded temporal periods in which an actual epileptic activity was detected. We treated the edges of the lateral grid as 1D strips in order not to lose these data points. CSDs of the 4 electrodes residing in the corners of the lateral grid were not calculated since they had no neighboring electrodes on both sides, and similarly for the edges of the lateral strip. After CSD calculation, the signals went through the onset latency estimation procedure described in 2.7.3.

### Gamma-band power calculation

We calculated broadband gamma-band power by high-pass filtering the signals, with 30Hz cutoff 4^th^ order Butterworth causal filter, and then taking the absolute value of the Hilbert transform and squaring. Importantly, since we were interested in onset latency estimation, we used a causal filter to avoid artificial shift of the onset backward in time. The onset latency of the gamma signal was estimated with the same procedure described in 2.7.3.

## Results

### Onset latencies across the posterior cortex

Significant responses and corresponding onset-latency-estimates (OLEs) were found in 69 out of 112 electrodes (Fig. 2 and supporting information video S1). Seven of the 69 electrodes were marked as showing epileptic activity and 3 more were labeled as ‘bad’ due to other electrical artifacts (see methods) and were ignored in the ensuing report of the results. The full list of OLEs is reported in the supporting information (tables S1.1 and S1.2). Generally, the spatio-temporal distribution of the OLEs followed the expected longer latency with higher hierarchy rank. The earliest OLEs obtained were those of posterior occipital electrodes, around 50-60ms (but see also section ‘Very early visual response at posterior intra-parietal sulcus’), with relatively little inter-trial variability (see Fig. 4 for single trials): 50 ± 4, 60 ± 2, 60 ± 3.5 and 96 ± 3.5 ms (see section ‘The early parietal response is not due to volume conduction’ for an earlier OLE of the latter electrode using the CSD reference, which emphasizes local activity), in 4 electrodes located over V1 and V2; (see table 1 for the full list of OLE obtained). All reported OLEs were calculated with significance level of 0.02 (see methods). However, the results are not affected dramatically by varying the significance level of the threshold (Fig. S2).

**Figure 3.**
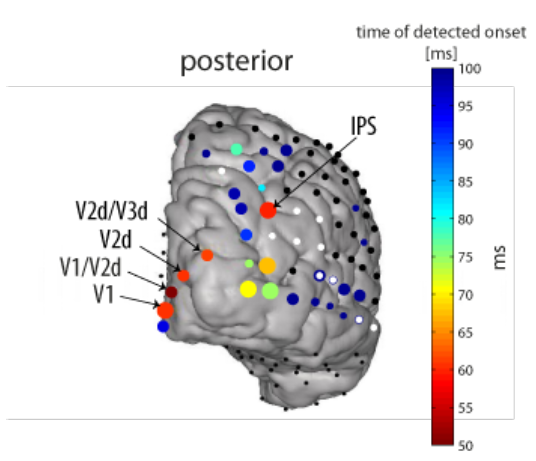
Posterior view of early onset latency estimates. The same as in Fig. 2, but with magnified temporal-scale, see color bar. Note that onset latencies estimated later than 100 ms are colored with the darkest blue on the scale. Labels according to retinotopic mapping as in Fig. 5 (see methods). Size of the circles is due to temporal error estimate, as in Fig. 2.

**Figure 4.**
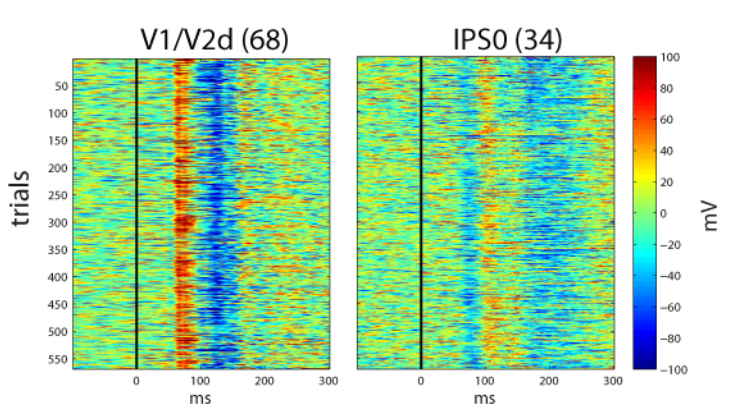
Event-related single trials in early responding electrodes. Electrode 68 (left) and electrode 34 (right). Electrodes are labeled from retinotopic mapping (see methods), as in Fig. 3 and 4. Note the remarkably consistent onsets in individual trials.

**Table 1.** Summary of early OLEs at electrodes having retinotopic labels for Patient 1, using common average reference (CAR), current source density (CSD) or broadband gamma power. Numbers are OLE ± temporal error estimate for the OLE (see methods for further explanation), in milliseconds. Missing electrodes in CSD were excluded from the analysis because they reside on the edge of the occipital strip and therefore are not appropriate for CSD estimation. Two electrodes did not have a significant OLE using the broadband gamma power signals.

| *Reference used*<br>*Electrode*<br>*localization* | *CAR* | *CSD* | *Gamma-band*<br>*Power* |
| --- | --- | --- | --- |
| *V1/V2d* | $50 \pm 4$ | $43 \pm 3.5$ | $47 \pm 2$ |
| *V1* | $60 \pm 2$ | $60 \pm 2$ | $64 \pm 5.5$ |
| *V2d* | $60 \pm 3.5$ | $58 \pm 1.5$ | $67 \pm 3$ |
| *V1/V2v* | $95 \pm 3.5$ | $76 \pm 30$ | ** |
| *V2d/V3d* | $61 \pm 3$ | * | $67 \pm 2.5$ |
| *V3v/V2v* | $170 \pm 10$ | * | ** |
| *IPS0* | $59 \pm 2.5$ | $60 \pm 2.5$ | $103 \pm 8.5$ |

\* - Electrode excluded from CSD analysis
\*\* - Electrode did not show a significant OLE

We also detected responses and calculated OLEs at various dorsal-parietal and ventral-temporal electrodes, varying between 60 and 200 ms (Fig. 2, and see supporting information table S1.1 for the full list of significant onsets). Generally, the more anterior the electrodes were located, the later the OLEs. However, there were distinctive exceptions to this rule. Electrode 34, located over posterior intra-parietal sulcus (IPS) showed a very early response, 59 ± 2.5 ms, a result we return to in more detail in the next section. Another exception was electrode number 55, which had an OLE of 107 ± 6.5 ms, earlier than other electrodes located posterior to it (see Fig. 2, lateral and posterior views). A general observation was that electrodes located in the dorsal stream (posterior parietal) responded overall relatively early, within 100 ms, consistent with the view of early processing in the dorsal visual system (Bullier, 2001; Bar, 2006; Chen *et al.*, 2007; Snyder *et al.*, 2012). In contrast, with a few exceptions, most electrodes on the ventral temporal surface responded later, ~100 – 200 ms OLEs.

Previously, (Parvizi *et al.*, 2012), showed a causal and specific role of two electrodes (72, 77 on the middle and posterior fusiform, abbreviated m-fus and p-fus, respectively) in this patient in face perception, by converging results from ECoG, fMRI and electrical brain stimulation. We report here the OLEs that were obtained for these electrodes: 84 ± 4 ms and 89 ± 4 ms respectively (Fig. 2, supporting information table S1.1). These electrodes, (together with their neighbor electrode 73) showed the earliest and most reliable onsets among the surrounding electrodes (Fig. 2). However, electrode 77 was marked as ‘bad’ due to electric artifacts (Fig. 2 and supplementary table S1.1).

### Very early visual response at posterior intra-parietal sulcus (IPS)

Whereas the general pattern of onset latencies followed the notion of a hierarchy, a significant and robust very early response of 59 ± 2.5 ms was measured in one electrode (no. 34; Fig. 2, 4) over the right posterior intra-parietal sulcus (IPS) (MNI coordinates - 35.12, −78.9, 41.89). fMRI retinotopic mapping placed the electrode over area IPS0 (see supporting information Fig. S4A). We verified that this early visual onset is not due to activity originating from V3A, a relatively low-level visual area (see supporting information Fig. S5). This response onset was nearly simultaneous with the response onset at most striate and peri-striate electrodes that we measured (electrodes 67, 69, 70 which were marked V1, V2d, V2d/V3d, respectively, based on retinotopic mapping, Fig. 5 and supporting information Fig. S4B). The only earlier responding electrode was electrode number 68, one of the electrodes located over V1/V2d, responding as early as 50 ± 4 ms (Fig. 5). Notably, this early response in area V1/V2d is due to a small voltage deflection that was marginally significant (see waveform in Fig. 5, for an earlier OLE see also results using CSD in section ‘The early parietal response is not due to volume conduction’). However, the main voltage deflection of electrode 68 was around 57 ms, nearly simultaneously with other early responding occipital electrodes and the IPS. All electrodes located around electrode 34 had later (or no) OLEs and larger temporal errors (see Fig. 3), supporting the claim that the response is localized and specific to the location of electrode 34.

**Figure 5.**
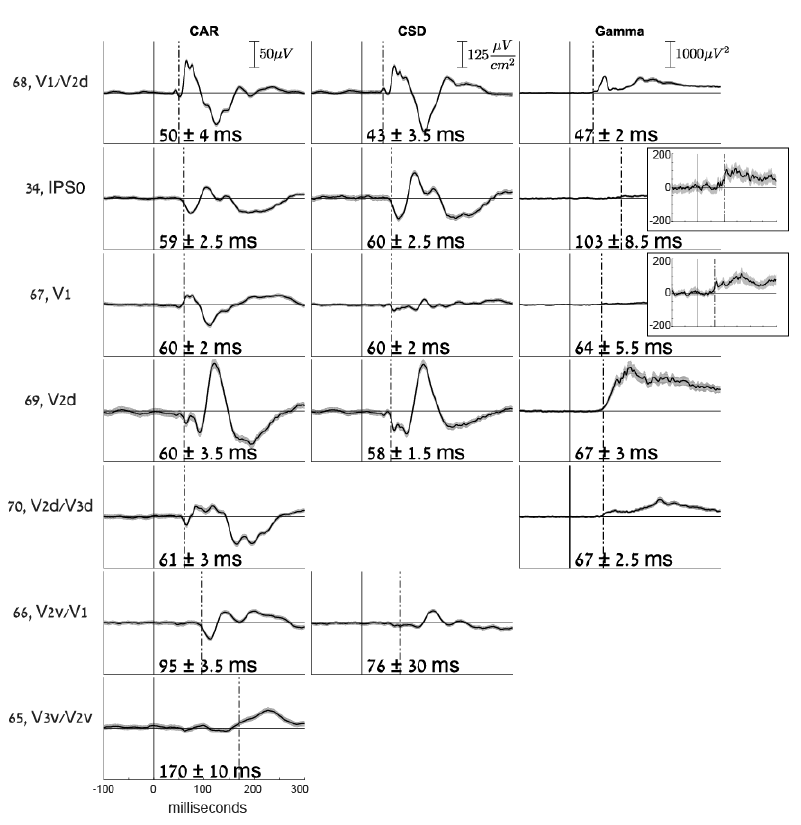
Waveforms and Onset Latency Estimates (OLEs). Comparing electrodes having retinotopic labels, using Common Average Reference (CAR), Current Source Density (CSD) or broadband gamma power (Gamma). To the left of each row, the number and retinotopic label of each electrode is specified. For each method, the scale is specified in the first row. Insets for broadband gamma power signals, electrodes 34 and 67, show a magnified y-scale in order to better view the responses. Missing electrodes in CSD were excluded from the analysis because they reside on the edge of the occipital strip and therefore are not appropriate for CSD estimation. Missing electrodes in the broadband gamma power case did not show a significant OLE (see Table 1)

Area MT/V5 is frequently noted in the literature as having a very early onset, comparable to and even earlier than V2 (ffytche *et al.*, 1995; Bullier, 2001), ascribed to pathways bypassing V1 (e.g. Sincich *et al.*, 2004; Nassi *et al.*, 2006). However, the early activation we report is unlikely to coincide with this area. In order to verify this, we localized area MT in the patient’s brain in three different ways. First, we converted the Talairach coordinates for MT, which were reported by Domoulin et al. (2000), to MNI space, using the nonlinear Yale MNI to Talairach coordinate conversion algorithm embedded in BioImage Suite 2.0, http://noodle.med.yale.edu/~papad/mni2tal/, based on Lacadie et al. (2008). We projected these MNI coordinates of MT onto the MNI-normalized brain of the patient. The presumed location of MT/V5 fell at the depth of a sulcus between electrodes 60 and 61 in the lateral grid. Second, an fMRI-based MT localizer was run on the same patient previously at Stanford Medical Center (unpublished data, courtesy of Dr. Corentin Jacques). Based on activation contrasts between radial grating motion versus static stimuli, the main MT activation was localized closest to electrodes 60-62 and 52-54 (Fig. S3A), very similar to the localization based on the coordinates reported by Domoulin et al. (2000). Third, we identified MT by retinotopic maps (Amano *et al.*, 2009), in particular the foveal representation that is distinct from the large foveal confluence of V1-V4. The results were very similar to the previous method (Fig. S3B). Unlike previous studies using non-invasive source estimation (ffytche *et al.*, 1995), we did not obtain very early onsets around the presumed location of MT, which is anatomically far from the IPS electrode 34 discussed above. The lack of early visual activity near the presumed MT location might be a consequence of the fact that MT was buried deep in a sulcus, far from any electrode, and due to the stationary stimuli, which were not optimal for activating this region. Notably, consistent with Sunaert et al., (1999), some motion-specific fMRI activation was observed using the motion localizer near the early responding electrode 34 at IPS (see supporting information Fig. S3A).

### The early parietal response is not due to volume conduction

Since the onsets of most early responding electrodes, including V1 and IPS, were almost simultaneous, it is possible that the IPS electrode actually captured volume-conducted activity from remote occipital electrodes. Several features of the response make this possibility unlikely. First, if the activity was volume conducted, electrodes in between the IPS and the medial occipital cortex should show this conducted activity as well (if not more strongly), but none of them, including other electrodes in the vicinity of IPS showed a similarly early response (see Fig. 3).

Second, we repeated the analysis of onset latency estimation in the same way as before, but using the current source density (CSD) event-related waveforms. As noted in the methods section ‘Current Source Density (CSD) estimation’, because each electrode is referenced to its neighbors, CSDs are much less sensitive to remote effects than are the CAR measures (Perrin *et al.*, 1987, and see Fig. 5; e.g. electrode 67 was attenuated in CSD vs. CAR, presumably due to the filtering out of remote influences). The results of this analysis are shown in Fig. 5 and Table 1. IPS electrode 34 still showed early activity (see Fig. 5) supporting a robust early localized response at posterior IPS.

One prominent difference in the OLEs obtained using CSD waveforms compared to CAR, was in the earliest OLE of medial occipital electrode 68 over V1/V2d; an early and small voltage deflection was now detected as significant and resulted in an even earlier OLE, at 43 ± 3.5 ms (see Fig. 5). Notably, this early response was significant but small, and still, the major response manifested in a much larger voltage increase was around 57 ms (Fig. 5).

### Latencies of broadband gamma power

The broadband gamma-band response (as opposed to the narrow-band oscillations elicited by specific stimuli) is considered as a correlate of local asynchronous neuronal firing (Manning *et al.*, 2009; Ray *et al.*, 2011; Miller *et al.*, 2014; Hermes *et al.*, 2017). We calculated the broadband gamma power (30 Hz high-pass cutoff, see methods) of all electrodes, and then performed the same onset latency estimation procedure on the power signals (see methods). Importantly, we used a causal filter in order not to shift voltage deflections backwards in time. The onsets detected in gamma power signals were generally not earlier than those detected using ERPs (see supporting information Table S1.3 for all obtained onsets). In the IPS electrode, the power seemed to start increasing around 70 ms post-onset, however, this increase only reached significance around 103 ms. See Table 1 and Fig. 5 for a comparison of onsets with retinotopic labels using the 3 methods (CAR; Common Average Reference, CSD; Current Source Density, or Gamma; Gamma-band power signals). See Supporting information Table S1.3 for the full list on OLEs.

## Discussion

The aim of this study was to demonstrate the spatiotemporal evolution of early visual information along the human posterior cortex. By calculating onset latencies using electrocorticographic surface electrodes located at occipital, posterior-temporal and parietal areas, we report upper bounds for the onsets of visual processing at various cortical sites. Generally, our results show a progression of latencies from early visual cortex along the ventral and dorsal streams of visual processing, as previously demonstrated in monkeys. However, we obtained a very early robust response over the posterior intra-parietal sulcus. This early response started around 60 ms, at about the same time that we measure responses over the earliest visual areas. Early access to the dorsal stream was postulated in recent theories of visual processing (Bullier, 2001; Bar, 2006; Chen *et al.*, 2007; Snyder *et al.*, 2012) but direct evidence for such a mechanism in humans is scarce.

According to the JuBrain Cytoarchitectonic atlas viewer (Mohlberg *et al.*, 2012, https://www.jubrain.fz-juelich.de/apps/cytoviewer/cytoviewer-main.php), using the MNI coordinates of the early IPS electrode, it is located in area PGp of the inferior parietal cortex, the most caudal area of human IPL. Congruently, the retinotopic mapping in our subject placed the electrode in area IPS0 (supporting information Fig. S4A). It is interesting to note that Caspers et al., (2013), found that area PGp contains receptor distributions very similar to those of ventral extrastriate visual cortex. Note however that since the electrode is located right over the inferior parietal sulcus in the native patient brain, it could capture as well activity originating from the ventral bank of the sulcus.

### Comparison to previous studies

In the monkey, several studies have reported visual onset latencies using invasive electrophysiological measurements. Bullier (2001) reviewed many such studies of single unit recordings in macaque monkeys. In this review, the latencies (medians reported first and then earliest 10% percentile in parenthesis) ranged between 45-80 ms (25-45) ms over V1 - e.g., 45 (25) ms in Maunsell and Gibson (1992), 55 (40) ms in Knierim and van Essen (1992), 85 (55) ms over V2 (Nowak *et al.*, 1995), 40-75 (45) ms over MT (Raiguel *et al.*, 1999) and 100 (70) ms over lateral intra-parietal (LIP) (Barash *et al.*, 1991). These latencies are generally longer than those we report here, which can be attributed to the fact that single unit spikes occur later than postsynaptic activity, which governs LFPs and ECoG signals.

A more recent study (Chen *et al.*, 2007) measured laminar field potentials in the macaque, which should be comparable to EEG and ECoG, since they measure average local activity that is correlated with pre-synaptic activity (Schroeder *et al.*, 1995; Chen *et al.*, 2007). These authors report 31 ms onset latencies over V1 and 33 ms over LIP. Using a rule of thumb of 3/5 ratio of conduction time between the monkey and human (Schroeder *et al.*, 2004; Chen *et al.*, 2007), our results compare well to these monkey studies. Ledberg *et al.* (2007), recording intracortical local field potentials, report longer onset latencies in striate cortex than Chen et al. (2007)(48, 55, and 67 ms in their 3 monkeys). Critically, while their earliest responses were always in the striate cortex, inferior parietal responses were seen within a few ms of the striate cortex, as early or earlier than peristriate cortex, and earlier than superior parietal, or inferior temporal cortex.

Direct evidence for onset latencies of visual responses in humans, with high spatial and temporal resolution, is scarce. In EEG, the C1 ERP component is considered to reflect V1 activity. This is due mainly to the fact that the C1 reverses polarity when the upper and lower visual fields are stimulated, consistent with the “cruciform” representation of these fields on the lower and upper banks of the calcarine sulcus (Clark *et al.*, 1994), although this association has been contested (Ales *et al.*, 2010). Based on the C1 scalp component and source estimation, (Foxe & Simpson, 2002) found an onset of 56 ms for a presumably V1 source and suggested that later activity was likely driven by a mixture of few more generators. Foxe and Simpson further report that by 70 ms activity is spread over dorso-parietal but not occipito-temporal cortex, which is only active by 80 ms. An earlier EEG study (ffytche *et al.*, 1995) using motion stimuli, found the earliest activation around 36 ms and more recent MEG studies suggested even earlier response in visual cortex. For example, Inui and colleagues (Inui & Ryusuke, 2006; Inui *et al.*, 2006) and Moradi (2003) using MEG, reported onset times earlier than 30 ms post onset in early visual cortex. Similarly, Shigihara and colleagues (Shigihara & Zeki, 2013; 2014; Shigihara *et al.*, 2016) find significant activity that peaks at around 30-40 ms but starts even earlier. Source reconstruction suggested that activity in this early period extended beyond V1. In one study (Shigihara *et al.*, 2016), activity was found to be significantly above baseline as early as 10-15 ms, although in this case the authors were more hesitant to ascribe this to cortical activity. Indeed, although MEG is arguably superior to EEG in the accuracy of source reconstruction, its resolution and accuracy is nevertheless constrained by the inverse problem.

Due to the proximity of electrodes to the generators, intracranial recordings offer high signal to noise and more precise localization (while providing a more limited spatial coverage). Kirchner et al. (2009) measured stereotactic intracranial EEG from an electrode on the superior bank of the calcarine sulcus in humans, and reported onsets around 25 ms. However, onsets were determined by eye rather than by any statistical measure. Yoshor et al. (2007), using grids and strips as in the current study, reported the earliest latencies around 56 ms post-onset, comparable to the current results, in electrodes presumably over V1-V2. However, they provide only rough localization of their electrodes relative to known visual areas. A rare multi-unit recording in humans from two electrodes in area V2-V3 reported onset latencies of just earlier than 60 ms (Self *et al.*, 2016).

Our results, with the advent of precise fMRI retinotopy of the same patient as well as precise electrode localization, show that for meaningful central images as used herein, activity in V1 starts earlier than 50 ms from stimulus onset, and that V2, as well as parietal sources, are also activated by 60 ms (table 1), and hence may contribute to the early VEP (e.g. to the C1 component).

It is hard to directly compare, or to make a strong statement, about the absolute latency of the earliest response in visual cortex. As noted, the numbers in humans range from as early as 10-15ms, to 60+ ms. This divergence very likely results from cardinal differences in the stimulation, the filtering parameters, and the statistical criteria for determining onsets. For example, data from non-human primates (Schroeder *et al.*, 1998) suggests that activation is faster for more peripheral than more central stimuli, and, most likely, larger stimuli would also produce earlier response simply by virtue of activating more neurons. Indeed, the results reviewed above showing very early activity in humans of 30 ms or earlier used stimuli like large checkerboard placed away from fixation, or even full field flashes. The luminance of the stimulus, and the degree of adaptation of retinal and cortical cells, is likely to affect the speed in which activation would be measureable as well. For example, the above mentioned studies by Inui et al. (Inui & Ryusuke, 2006; Inui *et al.*, 2006), which reported responses earlier than 30 ms used strong, full field flashes, after dark adapting their subjects for about 15 minutes. The content of the image may affect latencies as well, probably due to low level difference like spatial frequencies. For example, examining figure 7 in Kirchner et al.’s study (Kirchner *et al.*, 2009) suggests an earlier onset for checkerboard than for scenes in the human FEF. The method used to determine onset responses is also critical. Some studies use more lenient criteria, like eyeballing, to determine onsets, while others use more conservative and controlled methods like the permutation method used here, which would delay the determined onset. Thus, as noted before (Shigihara *et al.*, 2016), the relative activation latency of different areas within subject and paradigm seems to be of more interest than absolute latencies. In that sense, despite the different absolute latencies, our study is congruent with the previous conclusions from the MEG and EEG studies reviewed above, showing near simultaneous activation of striate and specific extra-striate cortex, as departure from a strictly hierarchical feedforward organization (cf. Zeki, 2016). Specifically, our results points to very early activity in the IPS.

Chambers et al. (2004), using TMS, showed that the right angular gyrus is involved in reorienting attention during two discrete temporal periods: early (90–120 ms) and late (210–240 ms) after target onset. Since our IPS electrode 34 was located right over the sulcus, it might reflect responses originating in both banks of the IPS. If the origin of the response is on the ventral bank of IPS (i.e., the angular gyrus), then the early IPS onset that we measured might be functionally related to the early period of attention orientation in Chambers et al.’s study, since the measured ERP waveform at the IPS indeed onsets at 59 ms but unfolds between 60 and 100 ms approximately, commensurate with Chambers et al.’s early period.

### Neuroanatomical origin of the early parietal response

How does visual stimulation reach the early IPS source so early? One possibility is that the IPS response is driven by direct projections from sub-cortical structures, bypassing V1. These V1-bypassing pathways are widely discussed in the context of ‘blindsight’ - a neuropsychological condition in which, despite damage to primary visual cortex, some visual abilities are preserved (Weiskrantz *et al.*, 1974; Barbur *et al.*, 1993). The sub-cortical structures projecting directly to extra-striate cortex can be either the thalamic lateral geniculate nucleus (LGN), or pulvinar nucleus (PN), reviewed recently by (Zeki, 2016). The other possibility is that very early responses in V1 are relayed directly to the IPS via feed-forward connections. Because we based our calculations on stimulus-locked averages, and the stimulus onset time was non-predictable, it is not probable that top-down connections drive this response directly (although top-down connections might alter sustained excitability and hence encourage earlier onsets). Next, we discuss these putative sources.

The LGN receives most of its input directly from parvo and magnocellular ganglion cells of the retina, although some inputs also pass via the superior colliculus (SC) (for a review, see Leopold, 2012). Projections from LGN to cortical areas other than V1 were demonstrated directly in the monkey. For example, Sincich et al. (2004) found a direct projection from LGN to area MT/V5 in the macaque using retrograde tracing. Others reported visually driven fMRI activation in several extastriate areas in V1-lesioned macaques, including V2,V3,V4, V5/MT, and lateral intraparietal area (LIP) (Schmid *et al.*, 2010). fMRI activation in all these sites, as well as behavioral detection of the visual target, was suppressed following reversible inactivation of the LGN. This provides a strong support for LGN being the source of the residual activation, although it does not determine whether the LGN was the direct source for any of the activated loci, nor it determines the latencies of the activity in the extrastriate regions. In humans, Bridge et al. (2008) showed evidence for an ipsilateral structural pathway between LGN and area MT / V5 in both blindsight patient and controls, using diffusion-weighted MRI. These findings support the existence of direct pathways from LGN to area MT/V5, other extrastriate regions, and perhaps parietal areas as well, bypassing V1. However, whether this pathway is fast enough to deliver signals into the IPS as early as 60 ms, is unknown. It is suggested that direct projections from LGN to the extrastriate and parietal cortex originate from koniocellular cells located at the intercalated layers of LGN (Hendry & Reid, 2000). Most koniocellular (K) projections are small and have thin axons (Reese & Guillery, 1987), and thus should have slower conduction velocities than the magnocellular projections to V1. This suggests that the koniocellular pathway is an unlikely candidate for explaining the fast parietal response we measured. However, some studies suggest that K cells are very heterogeneous morphologically and physiologically (Schiller & Malpeli, 1977; Hendry & Reid, 2000; Leopold, 2012).

Visual information can also enter the cortex bypassing V1 via the PN. The PN gets input through the retino-tectal pathway, via the SC. The retino-tectal pathway has long been thought of as linked functionally to the dorsal stream in regulating spatial attention, control of eye movements and visually driven behavior (Petersen *et al.*, 1987; Rafal *et al.*, 1991; Sapir *et al.*, 1999), which seem to require fast access of visual information. Indeed, it was lately established that a colliculo-pulvinar pathway projects to extrastriate cortex directly; Lyon et al., (2010) found a disynaptic pathway from SC to extra-striate cortex, passing through PN and from there to dorsal stream areas MT and V3 (but not to V2 or V4). Moreover, direct retinal afferent projections also innervate the same PN subdivision (inferior pulvinar nucleus, PIm), which projects to area MT, as recently found in the marmoset (Warner *et al.*, 2010; Leopold, 2012), and this pathway may provide fast cortical activation. Although there is no clear evidence for PN projections to parietal areas, considering the association of both the retino-tectal pathway and the inferior parietal cortex to spatial attention, this pathway may be involved in the fast responses we observed.

Finally, it is also possible that the early parietal response stems from the earliest responses measured in V1, which are relayed to the IPS via feed-forward connections. We measured a very early response at a single electrode located over V1/V2d (electrode 68), as early as 50 ± 4 ms using CAR, and even 43 ± 3.5 using CSD. Notably, this very early activity reflects a small voltage deflection in the ERP (see Fig. 5) while the following, and much larger response in the same electrode, as well as activity in all other striate and peri-striate electrodes onsets at approximately the same time as IPS, around 60 ms. It could still be the case that by the time activation in V1 and peri-striate areas becomes strong, the earliest weaker responses have already been carried forward and arrived to parietal cortex. Such mechanisms have been postulated before (VanRullen & Thorpe, 2002; Chen *et al.*, 2007), and the notion of ‘single spike wave propagation’ was even used for modeling of neural latency codes for vision that can account for fast visual performance (Thorpe *et al.*, 2001; VanRullen & Thorpe, 2002; Kirchner *et al.*, 2009). Importantly, this fast transfer of information from V1 to IPS in about 10-15 ms would require 1 or 2 synapses in between them. According to (Felleman & Van Essen, 1991), in macaques there is a direct connection between V1 and posterior parietal area (PIP) as well as from V2 to PIP, but not from V1-2 to lateral intra-parietal (LIP). To our knowledge, there is no direct evidence for such connections in humans.

### Very early visual processing in the dorsal stream

The early parietal response onset we report here is in accordance with current theories of visual information processing in the brain as well as with empirical evidence. For instance, Chen et al. (2007) show dorsal-to-ventral stream latency advantage at several stages of the processing hierarchy in monkeys. Considering the functional role of the dorsal stream in spatial vision (Mishkin *et al.*, 1983), and in attending to locations in space, these lateral connections could serve to prepare the ventral system for further processing at a given location. Several theoretical claims were raised stressing the need for early access of relatively crude information to the visual system, serving to guide and prepare it for more detailed and efficient later processing (Bullier, 2001; Bar, 2006; Chen *et al.*, 2007; Snyder *et al.*, 2012). Bullier (2001) reviews evidence for early visual responses in primate parietal and frontal cortex belonging to the dorsal visual stream, which he calls all-together ‘the fast brain’. He postulates the importance of these fast responses in interacting with and modulating lower order visual areas for integration of global-to-local visual information via feedback connections. Similarly, the ‘frame-and-fill’ model suggests that object recognition is achieved by gradually integrating fine details into an already established whole (Bar, 2006; Snyder *et al.*, 2012). According to this view, structures in the dorsal visual stream have an important role in initial parsing of the stimulus and passing it on to prefrontal and ventral areas, which then bit by bit process further specific details. Our results provide the first direct electrophysiological evidence for very early parietal, dorsal stream response in the human brain, which is essentially assumed in all the discussed models.

### Limitations and conclusion

The current results are based on a single subject. One of the limitations of using ECoG is that electrode type and location are determined based on clinical needs and it is therefore hard to find another patient having electrodes located over the exact same areas. Additionally, the electrode grids are located on the surface of the cortex, and cover about 10 functional columns, or roughly 2-5 × 10^5^ neurons. Consequently, some neural responses may be missed if they are too weak, farther away from the electrode, or if the spatial orientation of the responding neuronal population in the tissue under the electrode is such that dipoles cancel each other. Another reason for possibly missing earlier responses is the analysis itself. Our onset latency estimation analysis is designed to bound the type I error (false alarms). There is no assurance that the test is sensitive enough to detect the earliest responses if they are weak relative to the noise. It is also the case that the illumination of pixels on an LCD screen is gradual, and the exact luminance level at which retinal responses are elicited is undetermined. Further, our dependent measure was a change in the evoked potential and broadband power. It is possible that other measures like single unit spiking or denser electrode arrays could show shorter latencies. For all these reasons, the onset latencies reported here should be taken as upper bounds for the onset of visual processing in specific cortical areas of the human brain of a single patient. Interestingly, the power of the broadband (gamma) response emerged later than the evoked response, especially in the IPS electrode. These differences stress the functional distinction between these two types of signals, as has been observed previously in the spatial domain (Winawer *et al.*, 2013). Whereas the evoked potentials and the broadband response have been associated with synaptic input potentials and local neural spiking, respectively, the relationship between these two signals is yet to be elucidated.

Another limitation stems from the process of determining the precise location of the ECoG electrodes on the cortex anatomically. Our methods, like most other ECoG studies, depend on co-registration of CT image depicting the electrodes, with an anatomical MRI scan that was done prior to electrode placement. The alignment of these images is not trivial, both because of difference between the imaging modalities and the fact that the brain itself might move a bit during the surgery. While several methods have been developed in order to overcome these limitations (e.g. (Dykstra *et al.*, 2012); (Hermes *et al.*, 2010)), the localization of electrodes should be taken with some error margin. For an example, we cannot be sure whether electrode 34 was located more over the ventral or dorsal banks of the intra-parietal sulcus.

Nevertheless, considering the roughly 5:3 ratio between human and monkey latencies, respectively, due to size differences (Schroeder *et al.*, 2004; Chen *et al.*, 2007), the latencies we find are consistent with the reports from the monkey using penetrating electrodes and single unit recordings. Despite the discussed limitations, to our knowledge this is the first direct report of the onset timing of visual processing across multiple spatially localized sites in the human cortex, and specifically for the very early parietal response. Additional such measurements are needed for establishment of the generality of these results and for further exact characterization of the timing of early visual processing in the human cortex. Analysis schemes as devised here can easily be applied to existing bulks of ECoG data recorded for other purposes, for systematic examination of the onsets of processing in various cortical areas.

## Acknowledgements

The authors are grateful to Dr. Josef Parvizi for facilitating ECoG data collection from his patient and for fruitful discussions about the results. We acknowledge Vinitha Rangarajan and Bret Foster for assisting with data collection at the Stanford Medical Center. We thank Corentin Jacques for generously sharing with us the ECoG MT localizer data that he collected on the patient. We thank Israel Nelken for valuable consultations. This work was supported by grant 2013070 from the United-states Israel Binational Science Foundation to LYD and RTK and grant 1902_2014 from the Israel Science Foundation to LYD as well as NIH grant R01MH111417 to JW. TIR was supported by The Hoffman Leadership and Responsibility Fellowship Program at the Hebrew University of Jerusalem.

## Conflict of Interest Statement

The authors declare that they have no conflict of interest.

### Author Contributions

Conceptualization, L.Y.D. and T.I.R.; Methodology, T.I.R and L.Y.D.; Formal Analysis, T.I.R, J.W. and E.M.G; Resources, R.T.K.; Writing – Original Draft, T.I.R.; Writing – Review and Editing, L.Y.D., J.W. and R.T.K.; Visualization, T.I.R. and J.W.; Supervision, L.Y.D.; Funding Acquisition, L.Y.D., R.T.K and, J.W.

### Data Accessibility Statement

All analysis scripts are publicly available on <u>GitHub - https://github.com/tamaregev/visual-onset-latencies</u>. The data will soon be publicly available too. For any further requests, please contact the corresponding authors.

### Abbreviations

AG: – angular gyrus
CAR: – Common average reference
CSD: – Current source density
ECoG: – electrocorticography
ERP: – event related potential
IPS: – intraparietal sulcus
ISI: – inter-stimulus interval
K: – koniocellular
LGN: – lateral geniculate nucleus
LIP: – lateral intra-parietal
OLE: – Onset Latency estimate
PIP: – posterior parietal area
PN: – pulvinar nucleus
SC: – superior colliculus

## Supporting Information

### Movie S1 – Spatio-temporal evolution of visual responses along the posterior cortex (see electronic material)

The movie depicts the progression of significant visual responses in all electrodes over time, for 250 milliseconds. Three anatomical views are presented; posterior, inferior and medial. The electrode numbers and locations match those in Fig. 2 and Fig. S1. A green colored dot appears over an electrode whenever its activity exceeds the calculated threshold (see methods section ‘Onset latency estimation’ and Fig. 1). The colors represent the absolute amplitude of the response, such that lighter greens represent higher absolute amplitude.

#### 1. Full tables of all electrodes tested, significant onset latency estimates (OLEs) with their temporal errors

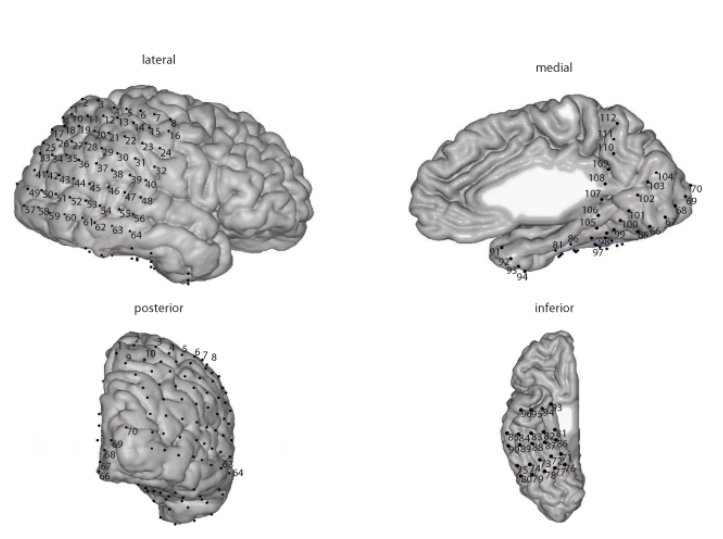
Electrode numbers in the following tables correspond to the figure above.

**Table S1.1.**
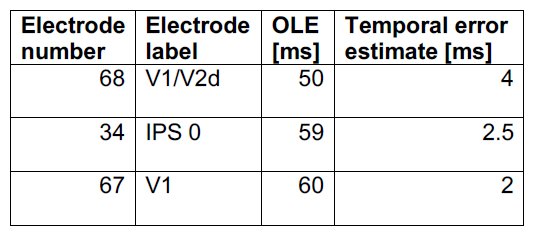
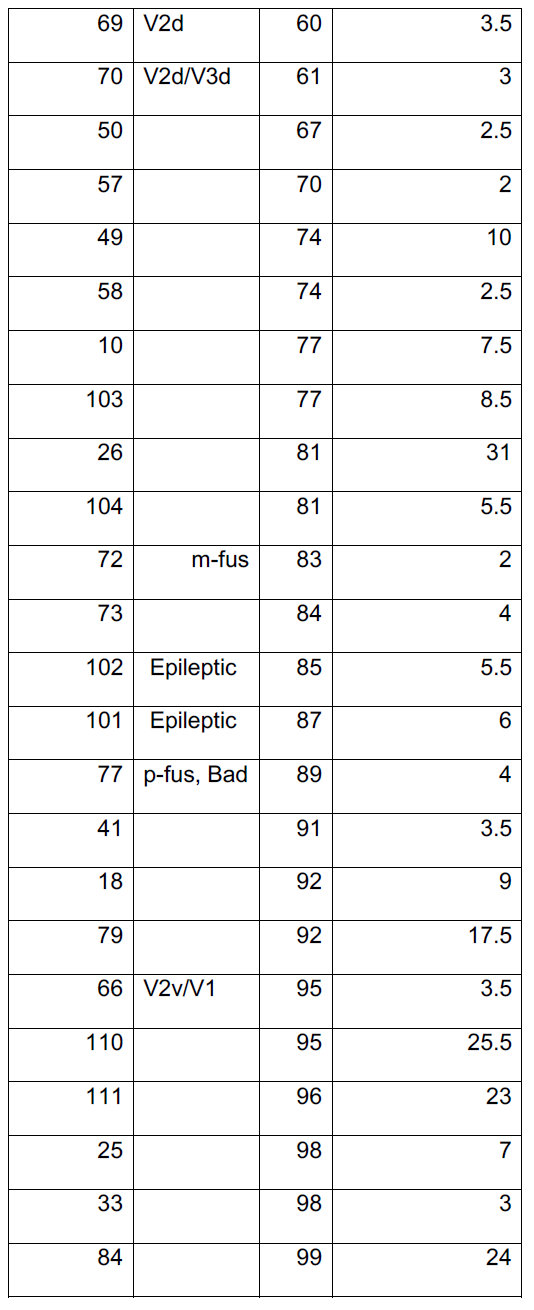
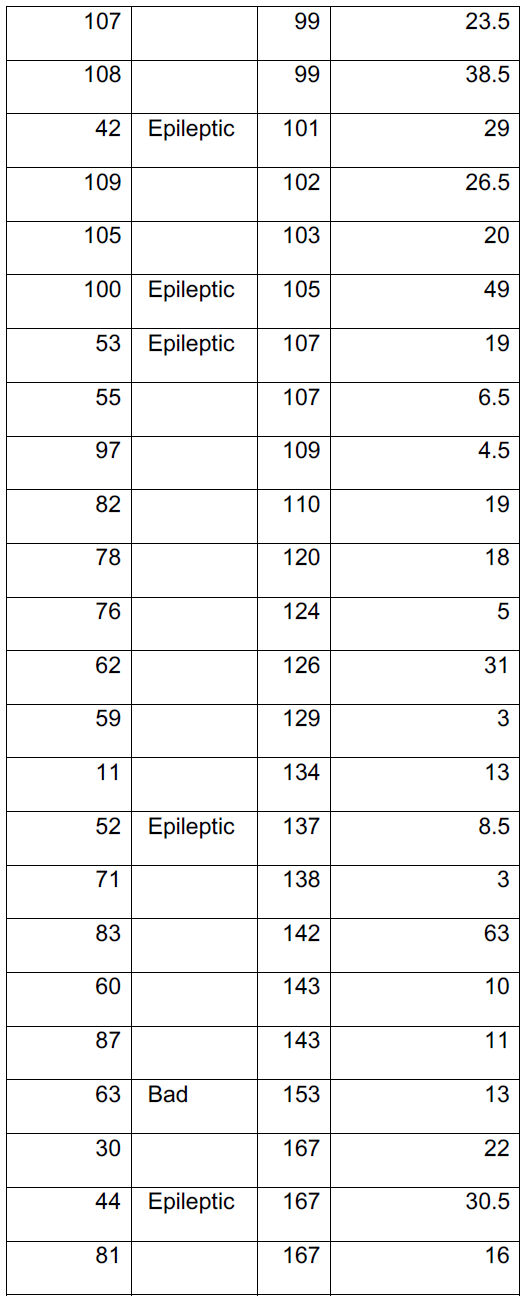
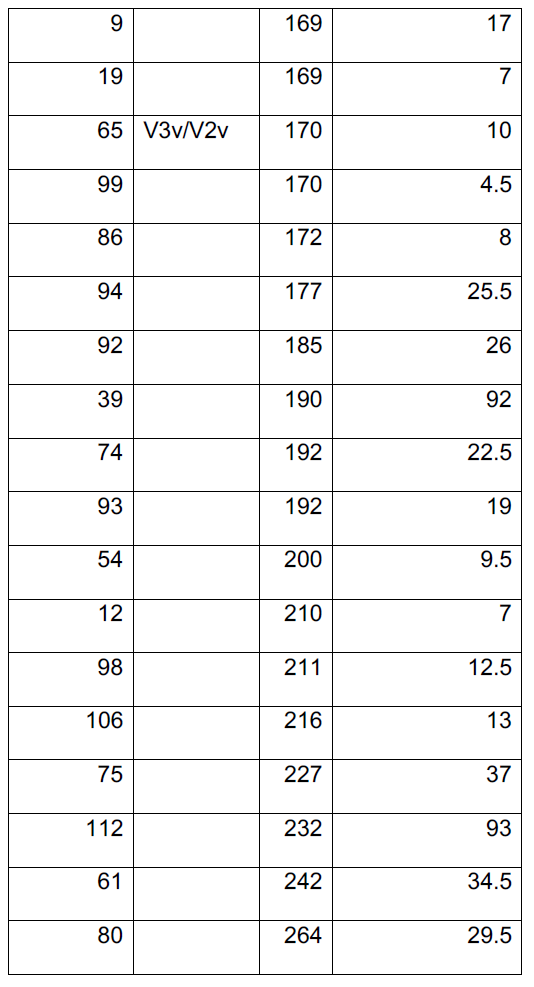
All significant OLEs, using common average reference (CAR) Out of 112 tested electrodes (numbers 1-112) 69 electrodes showed significant OLEs earlier than 300 ms, 7 of which were marked as epileptic and 2 more were labeled as ‘bad’ due to other electrical artifacts (see methods). Electrodes are ordered chronologically by OLE from early to late:

| Electrode number | Electrode label | OLE [ms] | Temporal error estimate [ms] |
| --- | --- | --- | --- |
| 68 | V1/V2d | 50 | 4 |
| 34 | IPS 0 | 59 | 2.5 |
| 67 | V1 | 60 | 2 |
| 69 | V2d | 60 | 3.5 |
| 70 | V2d/V3d | 61 | 3 |
| 50 |  | 67 | 2.5 |
| 57 |  | 70 | 2 |
| 49 |  | 74 | 10 |
| 58 |  | 74 | 2.5 |
| 10 |  | 77 | 7.5 |
| 103 |  | 77 | 8.5 |
| 26 |  | 81 | 31 |
| 104 |  | 81 | 5.5 |
| 72 | m-fus | 83 | 2 |
| 73 |  | 84 | 4 |
| 102 | Epileptic | 85 | 5.5 |
| 101 | Epileptic | 87 | 6 |
| 77 | p-fus, Bad | 89 | 4 |
| 41 |  | 91 | 3.5 |
| 18 |  | 92 | 9 |
| 79 |  | 92 | 17.5 |
| 66 | V2v/V1 | 95 | 3.5 |
| 110 |  | 95 | 25.5 |
| 111 |  | 96 | 23 |
| 25 |  | 98 | 7 |
| 33 |  | 98 | 3 |
| 84 |  | 99 | 24 |
| 107 |  | 99 | 23.5 |
| 108 |  | 99 | 38.5 |
| 42 | Epileptic | 101 | 29 |
| 109 |  | 102 | 26.5 |
| 105 |  | 103 | 20 |
| 100 | Epileptic | 105 | 49 |
| 53 | Epileptic | 107 | 19 |
| 55 |  | 107 | 6.5 |
| 97 |  | 109 | 4.5 |
| 82 |  | 110 | 19 |
| 78 |  | 120 | 18 |
| 76 |  | 124 | 5 |
| 62 |  | 126 | 31 |
| 59 |  | 129 | 3 |
| 11 |  | 134 | 13 |
| 52 | Epileptic | 137 | 8.5 |
| 71 |  | 138 | 3 |
| 83 |  | 142 | 63 |
| 60 |  | 143 | 10 |
| 87 |  | 143 | 11 |
| 63 | Bad | 153 | 13 |
| 30 |  | 167 | 22 |
| 44 | Epileptic | 167 | 30.5 |
| 81 |  | 167 | 16 |
| 9 |  | 169 | 17 |
| 19 |  | 169 | 7 |
| 65 | V3v/V2v | 170 | 10 |
| 99 |  | 170 | 4.5 |
| 86 |  | 172 | 8 |
| 94 |  | 177 | 25.5 |
| 92 |  | 185 | 26 |
| 39 |  | 190 | 92 |
| 74 |  | 192 | 22.5 |
| 93 |  | 192 | 19 |
| 54 |  | 200 | 9.5 |
| 12 |  | 210 | 7 |
| 98 |  | 211 | 12.5 |
| 106 |  | 216 | 13 |
| 75 |  | 227 | 37 |
| 112 |  | 232 | 93 |
| 61 |  | 242 | 34.5 |
| 80 |  | 264 | 29.5 |

**Table S1.2.**
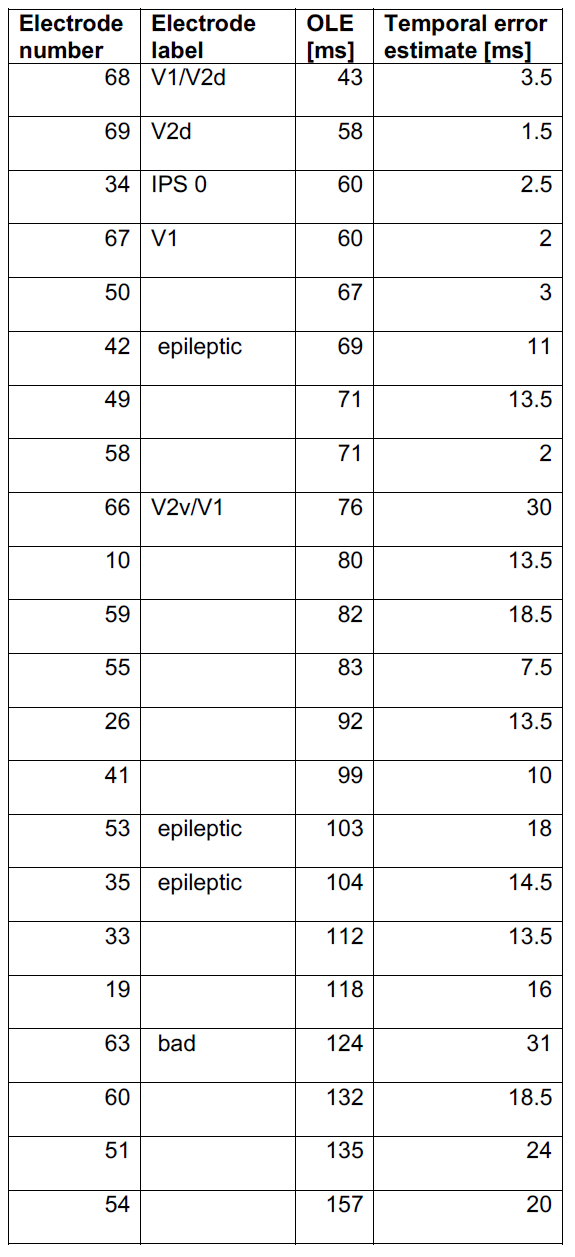
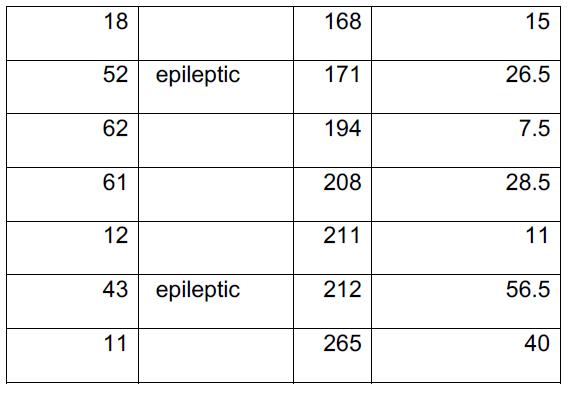
All significant OLEs, using current source density (CSD) reference. Out of 64 tested electrodes (lateral grid electrodes 2 to 63, excluding 8,57 because they reside in corners (see methods), and occipital strip 66 to 69), 29 electrodes showed significant OLEs earlier than 300 ms, 5 of which were marked as epileptic and 1 more was labeled as ‘bad’ due to other electrical artifacts. Electrodes are ordered chronologically by OLE from early to late:

**Table S1.3.** All significant OLEs, using broadband gamma power. Out of 112 tested electrodes (numbers 1-112), 25 electrodes showed significant OLEs earlier than 300 ms, 1 of which was marked as epileptic. Electrodes are ordered chronologically by OLE from early to late:

| **Electrode number** | **Electrode label** | **OLE [ms]** | **Temporal error estimate [ms]** |
| --- | --- | --- | --- |
| 68 | V1/V2d | 47 | 2 |
| 67 | V1 | 64 | 5.5 |
| 69 | V2d | 67 | 3 |
| 70 | V2d/V3d | 67 | 2.5 |
| 57 |  | 71 | 4 |
| 50 |  | 86 | 3.5 |
| 49 |  | 87 | 15 |
| 33 |  | 103 | 14 |
| 34 | IPS 0 | 103 | 8.5 |
| 72 | m-fus | 112 | 6.5 |
| 55 |  | 113 | 103.5 |
| 58 |  | 128 | 53.5 |
| 74 |  | 129 | 10 |
| 73 |  | 130 | 9 |
| 97 |  | 143 | 6.5 |
| 78 |  | 146 | 13 |
| 18 |  | 147 | 9 |
| 59 |  | 153 | 22.5 |
| 87 |  | 162 | 13 |
| 82 |  | 222 | 58.5 |
| 104 |  | 245 | 60.5 |
| 76 |  | 247 | 38.5 |
| 106 |  | 249 | 30.5 |
| 53 | epileptic | 274 | 74.5 |
| 81 |  | 303 | 11.5 |

#### 2. Several threshold analysis

**Fig. S2.**
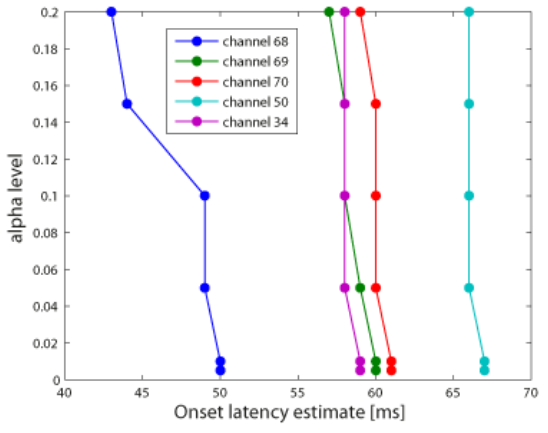
Onset latency estimations (OLEs) are not affected much by varying the alpha level chosen. We ran the bootstrapping procedure for calculating response thresholds (see methods) several times using various alpha levels – 0.005, 0.01, 0.05, 0.1, 0.15, and 0.2, using 5000 permutations (results reported in the paper are for alpha level of 0.01 and 4000 permutations). We used the common average referenced (CAR) signals. OLEs were affected only slightly by the varying alpha level, such that the OLE tended to be earlier, as expected, for larger alpha levels. This demonstrated well that our OLEs are an upper-limit. In electrode 68, the earliest responding electrode, the OLE was particularly reduced when using an alpha level greater than 0.1. This was due to an early and small voltage deflection, which was marginally significant. Interestingly, in the CSD analysis, this early response became significant.

#### 3. Motion localizer activation

**Figure S3A.**
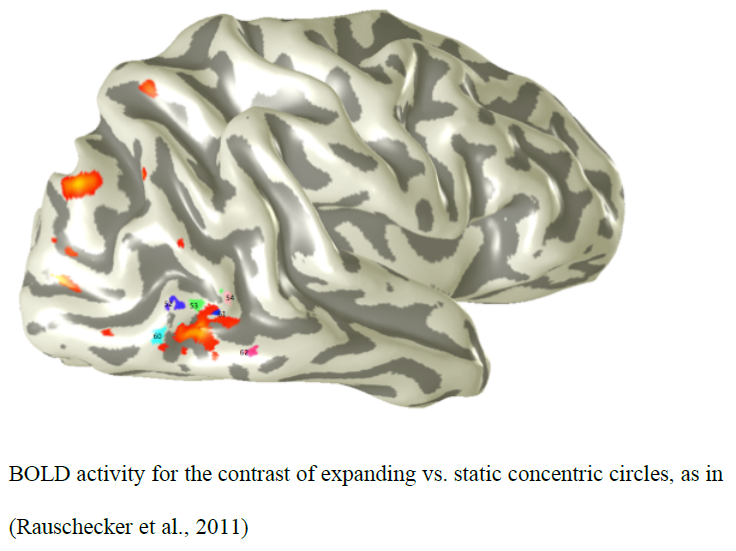
BOLD activation contrasting moving versus static stimuli. (Figure courtesy of Dr. Corentin Jacques)

**Figure S3B.**
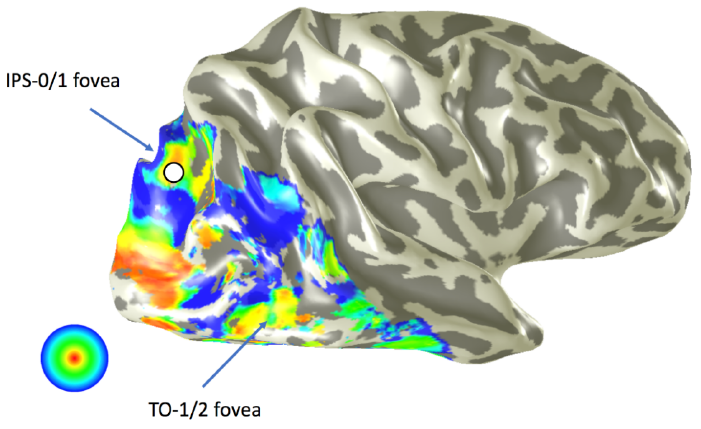
Retinotopic activations localizing area MT. (Amano et al, 2009).

#### 4. Visual maps - retinotopy

**Figure S4A.**
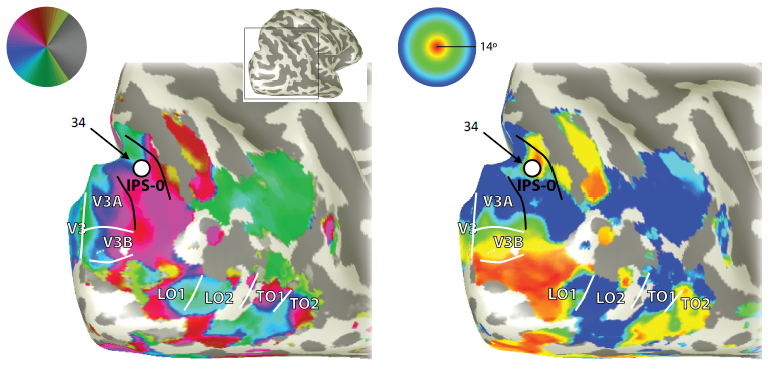
Electrode 34 over IPS0. The two main images show the smoothed (inflated) cortical surface of the right hemisphere, viewed from behind and the right side of the brain. The small inset indicates the location of the magnified views (black square). The color overlays show parameters from population receptive field model fits to a sweeping bar stimulus. Left - visual angle. Right – eccentricity. Electrode 34 (white disc with black outline) is located within the IPS-0 map, in the foveal confluence at the center of the IPS-0/1 cluster (Swisher et. al, 2007, Mackey, Winawer, Curtis, 2017). The diameter of the circle representing the electrode location is approximately 5 mm.

**Figure S4B.**
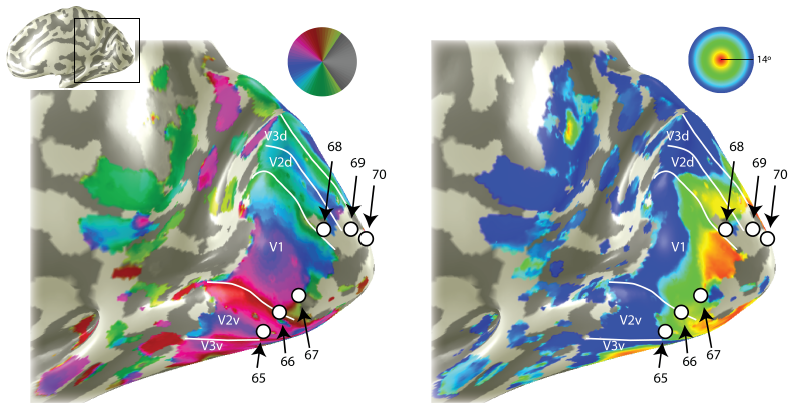
Occipital electrodes over V1, V2, V3. Visual maps as in Fig. S4A, but from the medial view. Left - visual angle. Right – eccentricity. Electrodes 65-70 belong to a single occipital strip. They appear to be unequally spaced due to smoothing (inflation) of the cortical surfaces.

#### 5. Electrode 34 is far from area V3-A

Since area V3-A is a relatively low-level visual area, we wanted to verify that the early visual onset measured in electrode 34, is not due to activity originating from area V3-A. For this reason, we marked areas V3-A, IPS-0 and IPS-1, as determined by retinotopic visual maps (see previous section), as well as electrode 34 on a volumetric brain image. As can be seen in Fig. S5, area IPS-0 (black) is closest to the electrode (red). Although the IPS-0 map borders V3-A, area V3-A (green) is far from electrode 34 (~1.5 cm). The electrode is increased in size for visibility, depicted as a 5-mm-diameter sphere. The center of the sphere is the location determined by the electrode localization procedure (see methods). We conclude that the early visual response in electrode 34 originates from area IPS-0.

**Figure S5.**
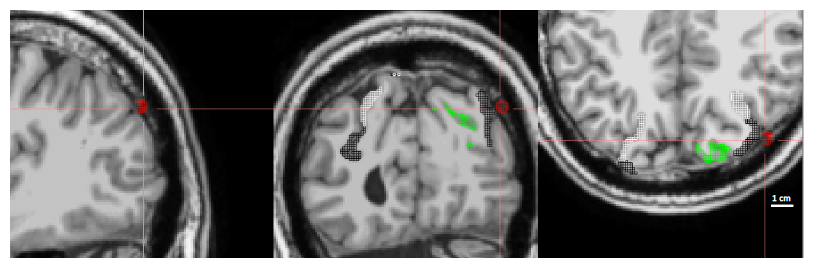
Volumetric localization of electrode 34. A volumetric image of the brain of patient 1. Red: electrode 34, black: IPS-0, white: IPS-1, Green: V3-A.

